# Genome Sequencing Identifies Previously Unrecognized *Klebsiella pneumoniae* Outbreaks in Neonatal Intensive Care Units in the Philippines

**DOI:** 10.1101/2021.06.22.449363

**Authors:** Celia C. Carlos, Melissa Ana L. Masim, Marietta L. Lagrada, June M. Gayeta, Polle Krystle V. Macaranas, Sonia B. Sia, Maria Adelina M. Facun, Janziel Fiel C. Palarca, Agnettah M. Olorosa, Gicell Anne C. Cueno, Monica Abrudan, Khalil Abudahab, Silvia Argimón, Mihir Kekre, Anthony Underwood, John Stelling, David M. Aanensen, the NIHR Global Health Research Unit on Genomic Surveillance of Antimicrobial Resistance

## Abstract

**Background:** *Klebsiella pneumoniae* is a critically important pathogen in the Philippines. Isolates are commonly resistant to at least two classes of antibiotics, yet mechanisms and spread of its resistance are not well studied.

**Methods:** A retrospective sequencing survey was performed on carbapenem-, extended spectrum beta-lactam- and cephalosporin-resistant *Klebsiella pneumoniae* isolated at 20 antimicrobial resistance (AMR) surveillance sentinel sites from 2015-2017. We characterized 259 isolates using biochemical methods, antimicrobial susceptibility testing, and whole genome sequencing (WGS). Known AMR mechanisms were identified. Potential outbreaks were investigated by detecting clusters from epidemiologic, phenotypic and genome-derived data.

**Results:** Prevalent AMR mechanisms detected include *bla*_CTX-M-15_ (76.8%) and *bla*_NDM-1_ (37.5%). An epidemic IncFII(Yp) plasmid carrying *bla*_NDM-1_ was also detected in 46 isolates from 6 sentinel sites and 14 different sequence types (ST). This plasmid was also identified as the main vehicle of carbapenem resistance in 2 previously unrecognized local outbreaks of ST348 and ST283 at 2 different sentinel sites. A third local outbreak of ST397 was also identified but without the IncFII(Yp) plasmid. Isolates in each outbreak site showed identical STs, K- and O-loci, and similar resistance profiles and AMR genes. All outbreak isolates were collected from blood of children aged <1.

**Conclusion:** WGS provided an in-depth understanding of the epidemiology of AMR in the Philippines, which was not possible with only phenotypic and epidemiologic data. The identification of three previously unrecognized *Klebsiella* outbreaks highlights the utility of WGS in outbreak detection, as well as its importance in public health and in implementing infection control programs.

**summary:** Whole genome sequencing identified three distinct previously unrecognized local outbreaks in a retrospective study in the Philippines, along with an epidemic plasmid carrying antimicrobial resistance genes, highlighting its importance in antimicrobial resistance surveillance, outbreak detection and infection control.

## INTRODUCTION

Antimicrobial resistance (AMR) is a serious threat to public health, as antimicrobial-resistant pathogens limit therapeutic options and result in increased morbidity and mortality [1]. AMR is also perceived as a threat to the achievement of the Sustainable Development Goals [2].

AMR surveillance has conventionally been performed by monitoring distribution of antimicrobial-resistant pathogens in the population through phenotypic methods such as antimicrobial susceptibility testing (AST), standard culture, and bacterial serotyping for identification and characterization [3,4]. WHONET is also widely used for surveillance data collection and analysis, while SaTScan integrated with WHONET allows early and broad detection of event clusters using retrospective or prospective algorithms and other flexible spatial and/or temporal scan parameters [5–7].

In the Philippines, priorities and methods for surveillance of AMR, hospital-acquired, and community-acquired infections are determined by the local health care facility’s infection control committee in coordination with the microbiology laboratory [8]. Hospital bacterial isolates, antibiograms, and clustering of patient groups within the hospital network are monitored and reviewed in a set time frame, and semi-annual infection rates and antibiograms are reported to clinicians and administrators [8]. Suspected outbreaks or the occurrence of uncharacteristically large numbers of cases are reported to the local office of the National Epidemiology Center – Department of Health for appropriate action [8, 9].

Bacterial typing based on phenotypes, however, fails to distinguish isolates that have the same resistance profiles (RP) and isolates belonging to closely related *Klebsiella* species [3, 10, 11]. The development of molecular methods such as pulsed-field gel electrophoresis and multi-locus sequence typing (MLST) enabled molecular detection of relatedness of isolates, but these are labor-intensive, time-consuming, expensive, and often of a low resolution insufficient for outbreak analysis because they assay variation at small proportions of the genome [12]. The decreasing costs of WGS can address these limitations by providing high-resolution subtypes of AMR pathogens [13, 14] and identifying AMR genes and their location on bacterial chromosomes or on plasmids [15, 16]. Through genome-wide analysis, a more granular picture of the status of AMR can potentially be determined by demonstrating mechanisms of AMR, transmission of AMR genes, and relatedness of strains [14–16]. *K. pneumoniae* is considered a microorganism of public health importance in the Philippines and is classified as critically important by the World Health Organization [17]. It is one of the leading causes of hospital-acquired infections, especially among the immunocompromised [18, 19]. Carbapenem-resistant *K. pneumoniae* was first isolated in the Philippines between 1992 and 1994 and has been molecularly characterized in recent years [15, 20]. A local outbreak of carbapenem-resistant ST340 and its possible transmission through an IncFII(Yp) plasmid with *bla*_NDM-1_, *rmtC*, and *sul1* was also identified through retrospective WGS survey, leading to the hospital’s review of infection control protocols and implementation of more stringent programs [15].

In this study, a retrospective sequencing survey was undertaken on carbapenem-, cephalosporin-resistant, and/or extended spectrum beta-lactamase (ESBL)-producing *K. pneumoniae* isolated from 2015-2017 to provide genomic context for local prospective surveillance. WGS analysis identified 3 potential local outbreaks among neonates, which were confirmed by corroborating epidemiological information, such as close isolation times and overlapping locations.

## METHODS

### Bacterial Isolates

From 2295 *Klebsiella* isolates collected from 2015-2017 and stored in the Antimicrobial Resistance Surveillance Resistance Reference Laboratory (ARSRL) biobank, 263 (11.5%) exhibited resistance to one or more of the following: carbapenems, third- and fourth-generation cephalosporins, and/or ESBLs. These were selected using metadata stored in WHONET. The isolates were retrieved and resuscitated on Tryptic Soy Broth and incubated overnight at 35°C for re-identification and re-testing of key phenotypic resistance (Supplementary Methods). Invasive specimens were prioritized when both invasive and non-invasive specimens representing identical combinations of RP, sentinel site, and year of collection were available.

### DNA Extraction and Whole-Genome Sequencing

Genomic DNA was isolated using nexttec™ 1-Step DNA Isolation Kit for Bacteria (nexttec Biotechnologie GmbH, 20N. 904) in accordance with manufacturer’s instructions. DNA was quantified using Quantifluor® dsDNA System (Promega, E2670) and Quantus™ Fluorometer (Promega, E6150), and then sent to the Wellcome Trust Sanger Institute for sequencing using Illumina® HiSeq2500 platform with 100- or 250-base paired-end reads. A total of 259 (98.5%) isolates passed quality control and were included in the study. Raw sequence data generated were deposited in the European Nucleotide Archive under the project accession PRJEB29738. Run accessions are provided in Microreact projects linked in the figure descriptions [21].

### Bioinformatics Analysis

The following analyses were performed using pipelines developed within the National Institute for Health Research Global Health Research Unit on Genomic Surveillance of AMR: QC, De novo assembly, mapping based SNP phylogeny, AMR, and MLST predictions [22–29]. Briefly, the sequences were assembled using SPAdes and identified using BactInspector, and contamination was detected using confindr [30–32]. Fastqc, multiqc, and qualifyr were utilized for quality control [27–29]. SNP-based phylogeny was generated by mapping reads to a reference sequence using BWA mem; variants were called and filtered using bcftools, and a maximum likelihood phylogeny was produced using IQTree [33–35].

Pathogenwatch was used to identify MLST, K-, and O-loci, virulence factors, and plasmid replicons [36]. AMR genes were predicted using ARIBA 2.14.4 in conjunction with the NCBI AMR acquired gene and PointFinder databases [37–38]. Sequences were analyzed using PlasmidFinder and mapped against *bla*_NDM_ plasmids p13ARS_MMH0112-3 and p14ARS_MMH0055-5 [39]. Only those with ≥95% coverage were considered as matched. Results were collated and uploaded to Microreact for visualization [21].

### Outbreak Analysis

Isolates with identical locations, forming clusters in the phylogenetic tree, were inspected as potential outbreaks. Maximum-likelihood phylogenetic trees were generated for each cluster using reference genomes EuSCAPE_IL028, EuSCAPE_DK005, and SRR5514218. Epidemiologic data, AST results, and genotypic characteristics of isolates in each cluster were then investigated. Infection origin was computed based on date of admission and sample collection date. Positive isolates collected more than 2 days after hospital admission were determined to be hospital-acquired. Meanwhile, positive isolates collected 0-2 days before hospital admission were classified as community-acquired. WHONET-SaTSCan’s space-time scan permutation simulated prospective was utilized to look for statistical clusters among isolates recovered from the same site in 2015-2017 and characterized by the same RPs as the outbreak isolates [7]. SaTScan analysis was also extended to 2019 to check the persistence of the RP at the sentinel site. A maximum cluster length of 365 days and a recurrence interval (RI) of >365 days were set to exclude random signals that occur by chance alone and are of limited epidemiologic significance [40].

## RESULTS

### Isolate Distribution and Characteristics

The 259 *Klebsiella* isolates were collected between 2015 and 2017 by 20 of 26 Antimicrobial Resistance Surveillance Program sentinel sites representing 16 of 17 regions (Supplementary Table 1). Isolates were submitted as carbapenem-resistant (n=81, 31.2%), carbapenem- and cephalosporin-resistant (n=58, 22.4%), cephalosporin-resistant (n=11, 4.2%), ESBL-producing (n=62, 23.9%), and ESBL-producing and cephalosporin-resistant (n=47, 18.1%) (Supplementary Table 2).

Invasive isolates from blood (n=240, 92.7%) and CSF (n=13, 5.0%) were prioritized for WGS. A few non-invasive samples, i.e. urine (n=3), sputum (n=1), tracheal aspirate (n=1), and umbilical cord (n=1), were also analyzed based on their resistance profiles.

The majority of the isolates were from inpatients (n=253, 97.7%) and from hospital-acquired infections (n=171, 66.0%). Isolates were collected from patients aged <1 to 93 years old, but most were from patients <1 year old (n=145, 56.0%), composed of 84.8% neonates (0-28d) and 15.2% infants (29d-11mo). Hence, infections were mostly detected in the neonatal department (n=77, 29.7%). Other patients aged <1 were also admitted to the intensive care unit (ICU), pediatric, pediatric-ICU, mixed ward, and emergency departments.

### Species Identification and Sequence Type

In silico species identification resulted in 214 (82.6%) *Klebsiella pneumoniae*, 36 (13.9%) *Klebsiella quasipneumoniae* subsp. *similipneumoniae*, and 9 (3.5%) *Klebsiella quasipneumoniae* subsp. *quasipneumoniae* (Figure 1). The 45 *K. quasipneumoniae* were also correlated with 45 identified *bla*_OKP_ genes, a *K. quasipneumoniae* chromosomal marker encoding ampicillin resistance [11]. The overlapping biochemical phenotypes and lack of a stable classifier among these closely related species may account for the inability of conventional laboratory techniques to definitively differentiate them [11, 41].

**Figure 1.**
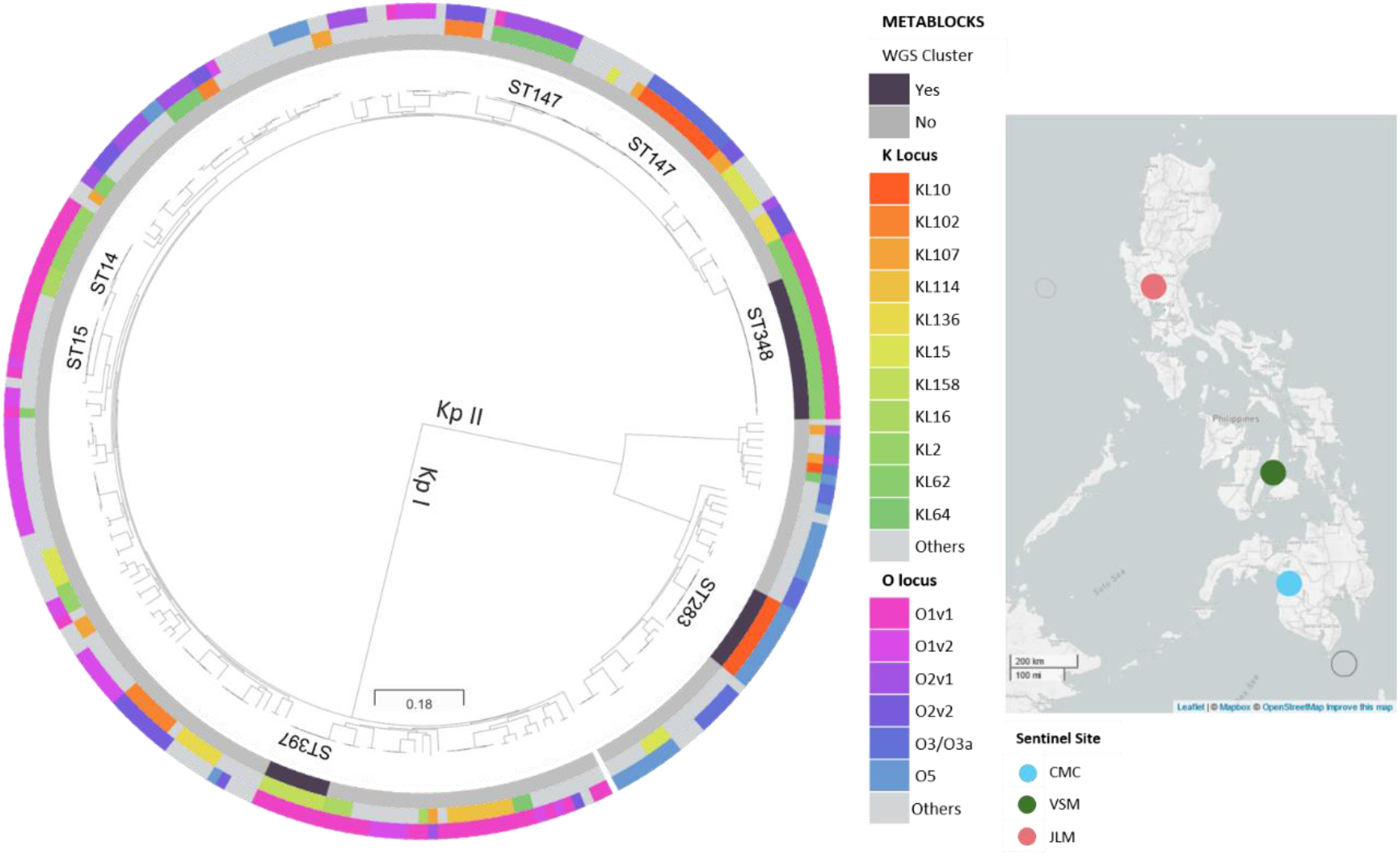
Phylogenetic tree of 259 *Klebsiella* isolates showing deep branches separating Kp I (*K. pneumoniae*) and Kp II (*K. quasipneumoniae*). Clusters based on linked genotypic (ST, KL, and O locus types) data showed 3 clusters of possible NICU outbreaks in 3 separate hospitals. Most ST348 isolates were collected in CMC, while ST397 and ST283 were unique in VSM and JLM respectively. Maximum-likelihood tree was inferred from mapping genomes to reference *K. pneumoniae* strain K2044 (GCA_009497695.1). This interactive view is available at: https://microreact.org/project/p8oycZe8jyu3Aghc3EE99c.

There were 102 different MLSTs predicted using ARIBA (Supplementary Figure 1). The most common were ST147 (n=23, 8.9%), ST348 (n=19, 7.3%), ST14 (n=10, 3.9%), and ST283 (n=9, 3.5%) (Supplementary Table 3). Ten isolates were identified to have novel ST profiles.

### Capsular and Lipopolysaccharide Typing

The *K. pneumoniae* capsule has been shown to be a key virulence determinant, suppressing host inflammatory response and providing resistance to antimicrobial peptides [42–44]. Capsular and lipopolysaccharide typing using the K and O loci identified 57 different K loci (KL) with good or higher confidence for 244 isolates, and 14 O loci for 256 isolates (Supplementary Figure 2). The most prevalent KL types were KL62 (n=22, 8.5%), KL10 (n=20, 7.7%), and KL64 (n=16, 6.2%). The most prevalent O loci were O1v1 (n=69, 26.6%), O1v2 (n=34, 13.1%), and O5 (n=31, 12.0%).

### Resistance Profiles and AMR Genes

A total of 93 RPs were observed, with the 4 most common accounting for 32.3% of the isolates (Supplementary Figure 1). RP-1, RP-3, and RP-4 were extremely drug resistant (XDR) RPs of *K. pneumoniae* and *Escherichia coli* reported to be expanding in the Philippines [15].

Known AMR mechanisms for carbapenem resistance, cephalosporin resistance, and ESBLs were identified (Supplementary Table 2). The most prevalent were the ESBL *bla*_CTX-M-15_ gene (n=199, 76.8%) and the carbapenemase *bla*_NDM-1_ gene (n=97, 37.5%). Other carbapenemase genes included *bla*_NDM-7_ (n=18, 6.9%), *bla*_KPC-2_ (n=4, 1.5%), and *bla*_NDM-9_ (n=1, 0.4%). Nineteen isolates had unknown mechanisms but carried combinations of *bla* genes and *ompK36* and *ompK37* mutations, which have been linked to carbapenem resistance in *Enterobacteriaceae* [45].

*rmtC*, encoding high resistance to aminoglycosides, was also identified in 48 (18.9%) isolates [46]. Other prevalent aminoglycoside resistance genes were: *aac(6’)-30/aac(6’)-lb’* (81.5%), *aac(6’)-Ib*(81.5%), *aac(3)-II* (46.3%), *aph(3”)-Ib* (47.9%), and *aph(6)-Id* (47.9%).

Fluoroquinolone resistance genes *oqxA* (95.4%) and *oqxB* (95.4%) were also observed in most isolates, along with *qnrB1* (32.8%), *qnrS1* (33.2%), *qnrB6* (12.7%), and *qnrB4* (4.2%). However, these genes only confer low level resistance or reduced susceptibility, which may not necessarily translate to phenotypic resistance [43]. Single *gyrA* mutations at codon 83 (11.2%) and double mutations at codon 83 and 87 (9.3%) were also found in some isolates. A single *parC* mutation co-occurred with 52 of 53 *gyrA* mutations (98.1%). *gyrA* and *parC* mutations corresponded to ciprofloxacin resistance in 50 of 53 isolates (94.3%). Sulfonamide and trimethoprim resistance genes *dfrA* (88.4%), *sul1* (70.7%), and *sul2* (58.3%) were also detected.

### Inc Type Profiling

A total of 32 different Inc types were observed among the 259 isolates (Supplementary Figure 1). IncFIB(K) (n=201, 21.4%) and IncFII(K) (n=163, 17.3%) were the most prevalent.

Notable Inc types identified were IncFII(Yp) (n=48, 5.8%) and IncX3 (n=21, 2.2%). A plasmid carrying IncFII(Yp), *bla*_NDM-1_, *rmtC*, and *sul1* (p13ARS_MMH0112-3) was previously linked to a nosocomial outbreak of *K. pneumoniae* ST340 in the Philippines [15]. The same IncFII(Yp) and AMR genes were also present in 46 of 48 IncFII(Yp)-positive isolates from 6 sentinel sites and 14 different STs (Supplementary Table 4). Short reads of all 46 isolates and one IncFII(Yp)-*bla*_NDM-7_ mapped to the plasmid with >95.0%coverage of the sequence length. There were 21 IncX3 replicons found in the genomes of 19 out of 20 *bla*_NDM-7_-carrying isolates from 9 sentinel sites and 8 STs (Supplementary Table 4). Short reads of all 19 were mapped to the *bla*_NDM-7_-carrying plasmid p14ARS_MMH0055-5 with 100% coverage of the plasmid sequence [15]. These results suggest that both IncFII(Yp) and IncX3 plasmids have been widely circulating and conferring carbapenemase resistance to a diverse genetic background, including non-epidemic strains, in the Philippines [15]. This study is limited by short read data, hence further plasmid or long-read sequencing will be conducted to characterize other vehicles of AMR gene transmission.

### Local Outbreak of *K. pneumoniae* ST348

WGS paired with epidemiological data provided a phylogenetic tree with 3 observed clusters (CMC, VSM, JLM) in 3 separate hospitals (Figure 1). This prompted an investigation to identify possible disease outbreaks. In all 3 clusters, all isolates were collected from blood and affected patients were all aged <1 year. Isolates in each cluster had identical MLST, capsular, and lipopolysaccharide types, and similar RPs and AMR genes (Figure 2). In addition, all the isolates in the CMC and JLM clusters carried plasmid p13ARS_MMH0112-3 (Supplementary Figure 3).

**Figure 2.**
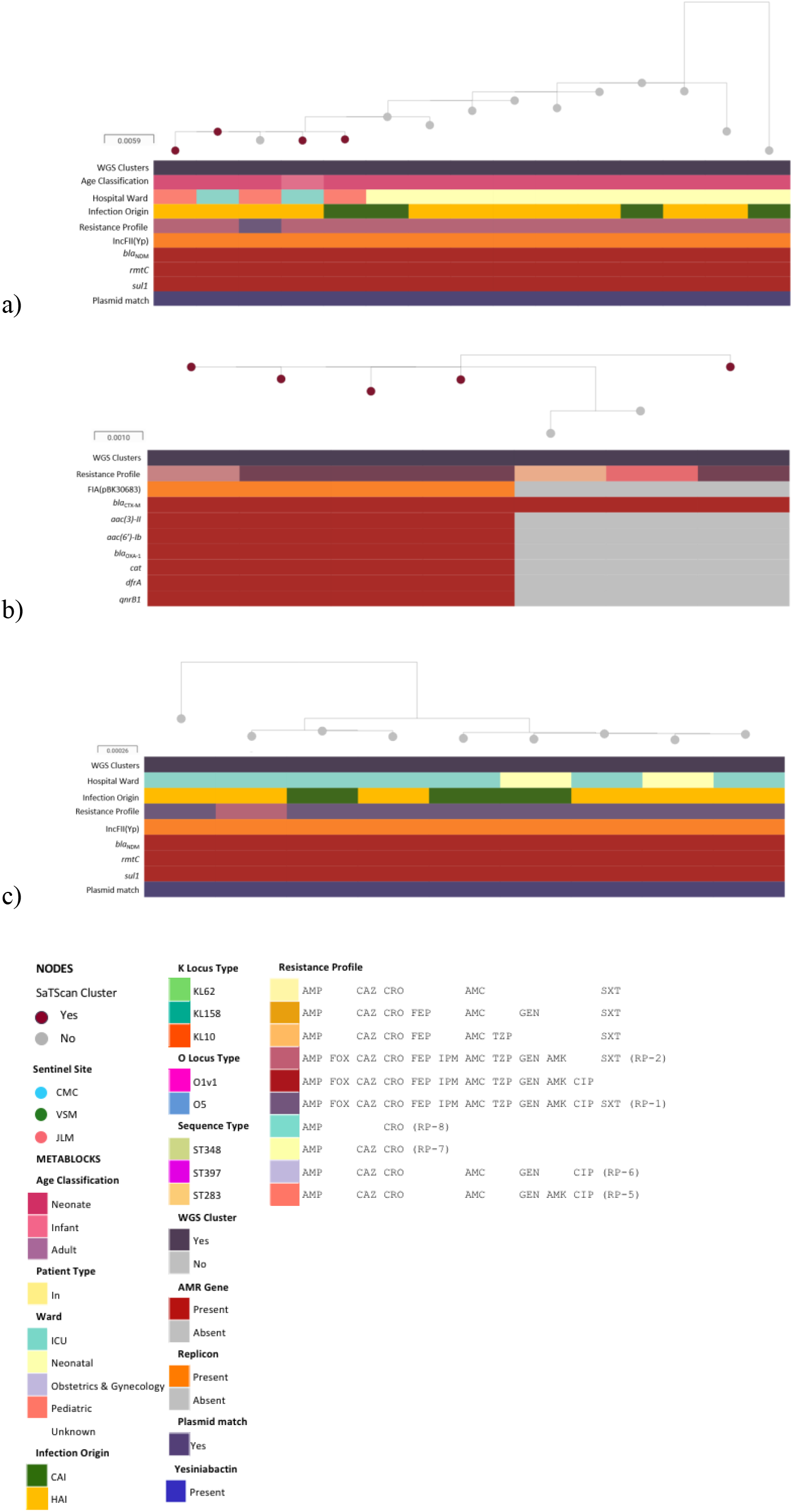
Phylogenetic tree, linked epidemiological and genotypic data of outbreak isolates at 3 sentinel sites. **A.** Maximum-likelihood tree of CMC ST348 (n=15) isolates was inferred from mapping genomes to reference EuSCAPE_IL028. This interactive view is available at: https://microreact.org/project/p5amCjPTePU6ggNWanXSe1/cb996623. **B.** Maximum-likelihood tree of 7 VSM ST397 isolates was inferred from mapping genomes to reference EuSCAPE_DK005. This interactive view is available at: https://microreact.org/project/9K27BJqkWxtpsqhovaK3B3/98829bfe. **C.** Maximum-likelihood tree of 9 JLM ST283 isolates was inferred from mapping genomes to reference SRR5514218. This interactive view is available at: https://microreact.org/project/7MrkhkdfoWuCQjiUuRscuS/075c339d. Infection origin was described as either hospital-acquired infection (HAI) or community-acquired infection (CAI). Presence or absence of the following AMR genes was also described: *bla*_NDM_, *rmtC, bla*_CTX-M_, *sul1, aac(3)-Il, aac(6’)-lb*, *bla*_OXA-1_, *cat, dfrA*, and *qnrB1*. Presence of plasmid replicons IncFII(Yp) and IncFIA(pBK30683) was also described, along with plasmid match (≥95% coverage) to p13ARS_MMH0112-3.

The largest cluster was observed in CMC, with 15 isolates collected between September 2016 and August 2017 identified as ST348 (Figure 2A). Most of the cases were hospital-acquired (n=11, 73.3%) with admissions to the neonatal ward (66.7%), pediatric ward (20.0%), and ICU (13.3%). All isolates had the same KL62 and O1v1 loci and carried virulence factor yersiniabactin (*ybt14*). Mean pairwise SNP difference between these 15 isolates was 6.3 SNPs (range: 1-28), suggesting intrahospital transmission when compared with the 53.1 mean pairwise SNP differences (range: 1-236) of other ST348 genomes from other sentinel sites that did not carry the *bla*_NDM_ plasmid p13ARS_MMH0112-3 [47, 48].

There were 14 ST348 isolates (93.3%) exhibiting RP-2, while 1 (6.7%) was additionally resistant to CIP exhibiting RP-1, which was confirmed by retesting the isolate (Figure 2A). Since *gyrA* and *parC* mutations were absent, this might have been caused by unknown mechanisms or differences in gene expression despite having similar low-level fluoroquinolone resistance genes, such as *oqx* or *aac(6’)-Ib-cr* [49].

### Local Outbreak of *K. pneumoniae* ST397

The second cluster was identified at sentinel site VSM, with 7 isolates collected from April 1 to May 2, 2016 identified as ST397. All patients were neonates, and cases were determined to be hospital-acquired (Figure 2B). KL158 and O1v1 loci and yersiniabactin gene *ybt9* were also identified in all isolates. The mean pairwise SNP difference between isolates was 3.9 SNPs (range: 1-8), indicating origin from a single transmission cluster [47, 48]. ST397 genomes from other hospitals were not available for comparison.

All isolates were ESBL-producing, with 57.1% (n=4) exhibiting RP-5 (full profile is shown in Figure 2), corresponding to isolates carrying IncFIA(pBK30683). Loss of IncFIA(pBK30683) in 3 isolates was also concordant with observed loss of *aac*, *bla*_OXA-1_, *cat, dfrA*, and *qnrB1*, indicating that these genes may be carried on an IncFIA(pBK30683) plasmid.

### Local Outbreak of *K. quasipneumoniae* ST283

The third cluster from JLM comprised 9 isolates collected from neonates in May 2016 to July 2017. Most cases were determined to be hospital-acquired (66.7%) (Figure 2C). All isolates were previously identified as *K. pneumoniae* by biochemical methods, but as *K. quasipneumoniae* subsp. *similipneumoniae* ST283 by WGS. All had the same KL10 and O5 loci. The mean pairwise SNP difference between isolates was 3.75 SNPs (range: 0-10), suggesting intrahospital outbreak [47, 48]. ST283 genomes from other hospitals were not available for comparison.

Eight isolates (88.9%) exhibited RP-1, while 1 (11.1%) additionally showed intermediate susceptibility to CIP (RP-2), which was confirmed by retesting the isolates. This is possibly due to unknown mechanisms or differences in gene expression, despite having the same *aac(6’)-Ib-cr, oqxAB*, and *qnrB1* genes conferring low level fluoroquinolone resistance, since *gyrA* and *parC* mutations were not detected [49].

### WHONET-SaTSCan Analysis

WHONET-SaTScan analysis of CMC *Klebsiella* isolates from 2015-2017 detected a statistically significant cluster for RP-2 (P<0.0013, RI=789d) (Table 1), which occurred in July 2017. This overlapped with the WGS-identified outbreak period in September 2016 to August 2017. Within this RP-2 cluster (n=10), we detected 3 of 4 ST37 isolates that mapped to the IncFII(Yp) plasmid (Supplementary Figure 3), and only 4 of 14 WGS-identified ST348 outbreak isolates (28.6%). Extending the analysis to 2019 also showed an RP-1 cluster (n=4) occurred in July to August 2018, indicating possible persistence of outbreak strain or of the epidemic plasmid.

**Table 1.** Clusters detected by WHONET-SaTScan analysis of CMC and VSM *Klebsiella* isolates from 2015-2019.

| RP No. | Resistance Profile | Recurrence Interval | Cluster p-value | Cluster Date | No. observed | No. of Outbreak isolates in cluster |
| --- | --- | --- | --- | --- | --- | --- |
| CMC |  |  |  |  |  |  |
| 1 | AMP FOX CAZ CRO FEP IPM AMC<br>TZP GEN AMK CIP SXT | 1,008 | 0.000992 | Jul-Aug 2018 | 4 | None |
| 2 | AMP FOX CAZ CRO FEP IPM AMC<br>TZP GEN AMK SXT | 789 | 0.00127 | Jul 2017 | 10 | 4 |
| VSM |  |  |  |  |  |  |
| 5 | AMP CAZ CRO AMC GEN AMK CIP | 10,369 | 0.0000964 | Apr 2016 | 4 | 4 |
| 6 | AMP CAZ CRO AMC GEN CIP | 4,575 | 0.000219 | Mar-Apr 2016 | 7 | 1 |
| 7 | AMP CAZ CRO |  | - | - | - | - |
| 8 | AMP CRO | 1,220 | 0.00082 | Dec 2018 | 3 | None |

Clusters were also detected in VSM, but only in 2 of 4 RPs (RP-5, RP-6) (Table 1). RP-6 (n=7) and RP-5 (n=4) clusters occurring in March-April 2016 and April 2016 respectively also fell within the WGS-identified outbreak period from April 1 to May 2, 2016. Both clusters were statistically significant (P<0.001), but the RP-5 cluster showed strongest statistical significance (P<0.000096, RI=10,369 days). Furthermore, only 5 of 7 WGS-identified ST397 outbreak isolates were detected in the WHONET-SaTScan clusters. Extending the analysis to 2019 also showed an RP-8 cluster (n=3) occurring in December 2018, indicating the possible persistence of AMR in VSM.

Lastly, WHONET-SaTSCan analysis of all JLM *Klebsiella* isolates (n=1,562) generated no clusters for either outbreak RP, although RP-1 and RP-2 have been observed in 12 and 29 cases respectively from 2015-2019 (Supplementary Figure 4). This suggests an even distribution or a gradual increase of cases, rather than a sudden increase, which is indicative of an outbreak signal detected by SaTScan. Altogether, results showed that cluster analysis with WHONET-SaTScan and a fixed set of parameters may not detect all clusters, especially if they are comprised of few isolates exhibiting more than one RP. Scan type and spatial and/or temporal parameters may, however, be refined to detect clusters not only among RPs but also in specific wards.

## DISCUSSION

We undertook a retrospective WGS survey of *Klebsiella pneumoniae* covering the years 2015-2017. We identified 3 outbreaks of *Klebsiella* among neonates in different hospitals in the Philippines, based on clusters observed in the phylogenetic tree, which resulted from combined epidemiologic and genotypic information. Average SNP differences in the 3 outbreaks were lower than the suggested thresholds of 16 and 21 SNPs for a *K. pneumoniae* intrahospital outbreak [47, 48].

Of the 3 outbreak strains, ST397 and ST283 were unique to VSM and JLM, respectively, but were nevertheless identified in other countries [50–53]. However, the same STs in other countries had differing AMR gene complements compared with the Philippine outbreak strains, which may therefore relate to local antimicrobial use practices [54].

For some isolates of the same outbreak strain, the complement of AMR genes differed among isolates of the same lineage, indicating distinct and potentially quite frequent gene-acquisition events, especially among hospital isolates. It is postulated that these AMR genes are acquired through selection due to antimicrobial exposure during hospital stay and may be carried by possibly epidemic plasmids circulating among non-epidemic strains, such as p13ARS_MMH0112-3, one of the main drivers of the JLM and CMC outbreaks [54]. Its occurrence in the genetic background of multiple STs within non-epidemic clones and at multiple sentinel sites suggests this is an epidemic plasmid that causes outbreaks among the immunocompromised, such as neonates in the ICU. However, further plasmid studies are needed to identify and characterize more AMR vehicles and modes of transmission.

There were no identified hypervirulent, extremely resistant *K. pneumoniae*, such as the epidemic KPC-producing ST258/ST11 clonal complex (CC258). However, possible XDR RPs (RP-1, RP-3, RP-4) (n=63, 24.2%) were among the most observed in this study, suggesting a possibly ongoing expansion [15]. As surveillance of AMR phenotypes is monitored, it may also be worthwhile to include surveillance of high-risk clones such as CC258 to predict invasive disease [54].

Using WGS, we were also able to distinguish what was phenotypically identified as *K. pneumoniae* to be *K. quasipneumoniae*. The phylogenetic tree of the 259 genomes in the Philippines showed deep branches separating the *K. pneumoniae* complex into *K. pneumoniae* and *K. quasipneumoniae*. *K. quasipneumoniae* is said to be less pathogenic than *K. pneumoniae*, which is more frequently associated with colonization or hospital-acquired infections [55].

The WGS pipeline was more efficient than WHONET-SaTScan at identifying potential outbreaks, since it was able to recognize the JLM outbreak, which SaTScan failed to do in a single run. However, WHONET-SaTScan may still be a good complement for WGS as demonstrated in CMC, since it was able to tag non-outbreak strains carrying the same epidemic plasmid as the confirmed outbreak isolates. Clinical interpretation is still based on the insights of clinical and infection control staff [7]. Further, the method may be limited by hospital policies governing the choice of antibiotics for testing and reporting of results, which impacts on the configuration of WHONET [56]. On the other hand, sequence data can provide a broader range of genotype information about isolates, which allows better characterization through improved molecular resolution.

Review of resistance profiles in the 3 hospitals showed persistence of the possible XDR RPs in CMC and JLM, indicating that more aggressive infection control interventions may be necessary to control the continuing expansion. We have communicated with the infection control staff of both hospitals to alert them, and they have implemented aggressive measures to prevent future outbreaks. These outbreaks illustrate that the routine utility of both WHONET-SaTScan and WGS in context of retrospective data will enable real-time generation of alerts, early outbreak detection and investigation, and immediate infection control [57].

In conclusion, WGS provided a more in-depth understanding of AMR epidemiology in the Philippines. This resulted in the identification of 3 previously unrecognized local outbreaks of *K. pneumoniae* and *K. quasipneumoniae* among neonates in 3 distinct areas, which was not possible using phenotypic data alone. Sustaining WGS can improve public health services to identify patients with distinct AMR sequences who are at risk of treatment failure, to predict potential outbreaks, and to take action for their immediate control.

## Acknowledgments

Members of the NIHR Global Health Research Unit on Genomic Surveillance of Antimicrobial Resistance: Harry Harste, Dawn Muddyman, Ben Taylor, Nicole Wheeler, and Sophia David of the Centre for Genomic Pathogen Surveillance, Big Data Institute, University of Oxford, Old Road Campus, Oxford, United Kingdom and Wellcome Genome Campus, Hinxton, UK; Pilar Donado-Godoy, Johan Fabian Bernal, Alejandra Arevalo, Maria Fernanda Valencia, and Erik C. D. Osma Castro of the Colombian Integrated Program for Antimicrobial Resistance Surveillance – Coipars, CI Tibaitatá, Corporación Colombiana de Investigación Agropecuaria (AGROSAVIA), Tibaitatá – Mosquera, Cundinamarca, Colombia; K. L. Ravikumar, Geetha Nagaraj, Varun Shamanna, Vandana Govindan, Akshata Prabhu, D. Sravani, M. R. Shincy, Steffimole Rose, and Ravishankar K.N of the Central Research Laboratory, Kempegowda Institute of Medical Sciences, Bengaluru, India; Iruka N Okeke, Anderson O. Oaikhena, Ayorinde O. Afolayan, Jolaade J Ajiboye, and Erkison Ewomazino Odih of the Department of Pharmaceutical Microbiology, Faculty of Pharmacy, University of Ibadan, Oyo State, Nigeria; Ali Molloy, alimolloy.com; and Carolin Vegvari, Imperial College London.

We are grateful to the members of the Antimicrobial Resistance Surveillance Program (Supplementary Note) that collected bacterial isolates and linked epidemiological data, especially to sentinel sites CMC, VSM, and JLM for cooperating and for providing additional information. We are also grateful to our project and support staff Elmer M. Herrera Jr., Laila T. Flores, Karis Lee D. Boehme, Michael F. Domingo, and our consultant Dr. Charmian M. Hufano for all their assistance.

