## Supplementary Information for "Genome Sequencing Identifies Previously Unrecognized *Klebsiella pneumoniae* Outbreaks in Neonatal Intensive Care Units in the Philippines"

[a] Members of the NIHR Global Health Research Unit for the Genomic Surveillance of Antimicrobial Resistance are listed in the Acknowledgments.

**Supplementary Method**

From 2015 to 2017, the Philippines' Department of Health Antimicrobial Resistance Surveillance Program (ARSP) had collected data from a total of 31,309 *K. pneumoniae* isolates representing 24 sentinel sites in 16 regions using the ARSP workflow [1]. Out of the collection, 2,295 (7.3%) isolates were referred to the Antimicrobial Resistance Surveillance Reference Laboratory (ARSRL) for confirmatory testing if they fall in one of the following categories: (1) resistant to carbapenems; (2) resistant to 3rd and 4th-generation cephalosporins collected on the first 10 days of the month; (3) ESBL-positive on the first 10 days of the month; and (4) *ampC*-positive on the first 10 days of the month. In 2017, isolates from the 4th category was replaced by all colistin-resistant isolates. Confirmatory testing in the ARSRL involved automated bacterial identification and antimicrobial susceptibility testing (AST) using VITEK® 2 Compact (bioMérieux) for the following antibiotics: ampicillin (AMP), ceftriaxone (CRO), ceftazidime (CAZ), cefepime (FEP), gentamicin (GEN), imipenem (IPM), ertapenem (ERT), amoxicillin-clavulanic acid (AMC), ciprofloxacin (CIP), and co-trimoxazole (SXT). Disk diffusion method was employed for amikacin 30ug (AMK), cefoxitin 30ug (FOX), meropenem 10ug (MEM), and piperacillin/tazobactam 100/10ug (TZP). ESBL production was confirmed by cefotaxime/cefotaxime with clavulanic acid (CT/CTL) (bioMérieux, 532240) or ceftazidime/ceftazidime with clavulanic acid (TZ/TZL) E-test MIC gradient strip (bioMérieux, 532540), whereas confirmation of carbapenemase production employed the Modified Hodge Test method. AST results were interpreted based on the interpretive criteria and breakpoints recommended by the latest version of the Clinical Laboratory Standards Institute [2]. All referred *K. pneumoniae* isolates were then lyophilized and stored in the ARSRL biobank facility.

#### SUPPLEMENTARY TABLES

**Supplementary Table 1.** Distribution of isolate sequence types and resistance profiles per sentinel site.

| Sentinel Site | No. of Isolates | No. of Observed ST | Prevalent ST (No. Observed) | No. of Observed RP | Prevalent RP (No. Observed) |
|---|---|---|---|---|---|
| BGH | 10 | 8 | 219 (2) | 8 | AMP CAZ CRO SXT (2) |
| BRH | 4 | 4 | | 4 | |
| BRT | 2 | 2 | | 2 | |
| CMC | 26 | 7 | 348 (15) | 6 | AMP FOX CAZ CRO FEP IPM AMC TZP GEN AMK SXT (RP-2) (20) |
| CRH | 3 | 2 | 20 (2) | 1 | AMP FOX CAZ CRO FEP IPM AMC TZP GEN AMK CIP SXT (RP-1) (3) |
| CVM | 28 | 13 | 70 (5) | 20 | RP-1 (7) |
| DMC | 8 | 8 | | 7 | AMP CAZ CRO AMC GEN SXT (2) |
| EVR | 20 | 14 | 15 (3) | 13 | AMP CAZ CRO FEP AMC GEN CIP SXT (4) |
| FEU | 2 | 2 | | 2 | |
| GMH | 8 | 4 | 36 (4) | | AMP CRO AMC SXT (3) |
| JLM | 33 | 20 | 283 (9) | 18 | RP-1 (12) |
| MAR | 6 | 5 | 147 (2) | 6 | |
| MMH | 19 | 12 | 852 (4) | 13 | AMP FOX CAZ CRO FEP IPM AMC TZP GEN AMK (4) |

| | | | | | |
|---|---|---|---|---|---|
| | | | 147 (4) | | |
| NKI | 6 | 6 | | 6 | |
| NMC | 19 | 16 | 14 (2)<br>147 (2)<br>1916 (2) | 14 | AMP FOX CAZ CRO FEP IPM AMC TZP GEN CIP SXT (RP-4) (3) |
| PGH | 9 | 8 | 307 (2) | 9 | |
| SLH | 1 | 1 | 1026 | 1 | AMP CAZ CRO FEP AMC --- SXT |
| STU | 9 | 4 | 147 (5) | 7 | AMP FOX CAZ CRO FEP IPM AMC TZP CIP SXT (RP-3) (3) |
| VSM | 43 | 25 | 397 (7) | 25 | RP-1(7) |
| ZMC | 3 | 3 | | 2 | AMP CAZ CRO FEP AMC TZP GEN AMK SXT (2) |

**Supplementary Table 2**. Distribution of isolates based on referral category and known resistance mechanisms.

| Resistance Mechanism | Carb | Carb, Ceph-R | Ceph-R | ESBL | ESBL, Ceph-R | **Total (%)** |
|---|---|---|---|---|---|---|
| *bla*$_{\text{NDM-1}}$ | 56 | 7 | | | | **63 (24.3%)** |
| *bla*$_{\text{NDM-7}}$ | 13 | | | | | **13 (5.0%)** |
| *bla*$_{\text{KPC-2}}$ | 1 | | | 1 | 1 | **2 (0.8%)** |
| *bla*$_{\text{CTX-M-3}}$ | | 1 | 1 | 5 | | **7 (2.7%)** |
| *bla*$_{\text{CTX-M-14}}$ | | | | | 2 | **2 (0.8%)** |

| | | | | | | |
|---|---|---|---|---|---|---|
| *bla*$_{CTX-M-15}$ | | 5 | 5 | 42 | 36 | **88 (34.0%)** |
| *bla*$_{NDM-1}$, *bla*$_{CTX-M-15}$ | | 32 | 1 | | | **33 (12.7%)** |
| *bla*$_{NDM-7}$, *bla*$_{CTX-M-15}$ | | 5 | | | | **5 (1.9%)** |
| *bla*$_{NDM-9}$, *bla*$_{CTX-M-15}$ | | 1 | | | | **1 (0.4%)** |
| *bla*$_{NDM-1}$, *bla*$_{CTX-M-15}$, *bla*$_{SHV-27}$ | | 1 | | | | **1 (0.4%)** |
| *bla*$_{KPC-2}$, *bla*$_{CTX-M-15}$ | | 1 | | | | **1 (0.4%)** |
| *bla*$_{CTX-M-14}$, *bla*$_{CTX-M-15}$ | | | | 1 | | **1 (0.4%)** |
| *bla*$_{SHV-12}$ | | | | | 1 | **1 (0.4%)** |
| *bla*$_{SHV-27}$ | | 1 | | | | **1 (0.4%)** |
| *bla*$_{SHV-28}$ | | | | 1 | 2 | **3 (1.2%)** |
| *bla*$_{CTX-M-3}$, *bla*$_{SHV-27}$ | | | | 1 | | **1 (0.4%)** |
| *bla*$_{CTX-M-3}$, *bla*$_{SHV-28}$ | | | | 3 | 1 | **4 (1.5%)** |
| *bla*$_{CTX-M-14}$, *bla*$_{SHV-27}$ | | | | 1 | 1 | **2 (0.8%)** |
| *bla*$_{CTX-M-15}$, *bla*$_{SHV-27}$ | | 1 | | | | **1 (0.4%)** |
| *bla*$_{CTX-M-15}$, *bla*$_{SHV-28}$ | | | | 7 | 3 | **10 (3.9%)** |
| Unknown | 11 | 3 | 4 | 1 | | **19 (7.3%)** |
| **Total (%)** | **81 (31.3%)** | **58 (22.4%)** | **11 (4.2%)** | **62 (23.9%)** | **47 (18.1%)** | **259 (100%)** |

**Supplementary Table 3.** Genotypic characteristics of the top 10 most prevalent STs.

| ST (No. observed) | O locus (No. observed) | K locus (No. observed) | Prevalent Inc Type Profile (No. observed) | Key AMR determinants based on referral category (No. observed) | | | | |
|---|---|---|---|---|---|---|---|---|
| | | | | **Carb R** | **Carb R, Ceph R** | **Ceph R** | **ESBL** | **ESBL, Ceph R** |
| 147 (23) | O2v1 (8) | KL64 (8) | Col440I; Col440I; IncFIB(pKPHS1); IncFIB(pQil); IncR; IncX3 (3) | *bla*$_{NDM-7}$ (5) | *bla*$_{NDM-1}$ (1) | | *bla*$_{CTX-M-15}$ (2) | |
| | O3/O3a (10) | KL10 (10) | Col440I; Col440I; FIA(pBK30683); IncA/C2; IncR (3)<br>Col440I; Col440I; FIA(pBK30683); IncFIB(K); IncFIB(Mar); IncFII(K); IncHI1B; IncR (2)<br>Col440I; FIA(pBK30683); IncR (2) | *bla*$_{NDM-7}$ (2)<br>*bla*$_{NDM-1}$ (6)<br>*bla*$_{CTX-M-15}$, *bla*$_{OXA-1}$, *bla*$_{SHV-11}$ (1) | | | *bla*$_{CTX-M-15}$ (1) | |
| | O3b (1) | KL14 (1) | IncFIB(pKPHS1); IncFII(K) | | | | | *bla*$_{CTX-M-15}$ (1) |
| | OL101 (3) | KL81 (3) | Col440I; IncFIB(K); IncFII(K); IncX3 (2) | *bla*$_{NDM-7}$ (2) | *bla*$_{NDM-9}$ (1) | | | |
| | OL104 (1) | - | Col440I; IncFIB(K); IncFII(K); IncX3 (1) | | | | *bla*$_{CTX-M-15}$ (1) | |

| | | | | | | | | |
|---|---|---|---|---|---|---|---|---|
| 348 (19) | O1v1 (19) | KL62 (19) | IncFIB(K)/IncFII(K); IncFIB(pB171); IncFIB(pKPHS1); IncFII(K); IncFII(Yp) (7)<br><br>IncFIB(K); IncFIB(pB171); IncFIB(pKPHS1); IncFII(K); IncFII(K); IncFII(Yp) (5) | *bla*$_{\text{NDM-1}}$ (4) | *bla*$_{\text{NDM-1}}$, *bla*$_{\text{CTX-M-15}}$ (12) | | *bla*$_{\text{CTX-M-15}}$ (2) | *bla*$_{\text{CTX-M-15}}$ (1) |
| 14 (10) | O1v1 (10) | KL2(7)<br>KL16(3) | Col440I; IncFIB(K); IncFII(K) (2)<br>IncN; IncU (2) | *bla*$_{\text{NDM-1}}$ (1) | | *bla*$_{\text{CTX-M-15}}$ (2)<br>*bla*$_{\text{CTX-M-3}}$(1)<br>*bla*$_{\text{SHV-28}}$ (1) | *bla*$_{\text{CTX-M-3}}$, *bla*$_{\text{SHV-28}}$ (2)<br>*bla*$_{\text{CTX-M-15}}$, *bla*$_{\text{SHV-28}}$ (1) | |
| 283 (9) | O5 (9) | KL10 (9) | Col440II; FIA(pBK30683); IncFIB(pB171); IncFII(Yp) (8) | *bla*$_{\text{NDM-1}}$ (8) | *bla*$_{\text{NDM-1}}$, *bla*$_{\text{CTX-M-15}}$ (1) | | | |
| 15 (8) | O1v1 (8) | KL112(6) | ColpVC; IncFIB(K); IncFIB(pKPHS1) (3)<br>ColpVC; IncFIB(K) (2) | | | *bla*$_{\text{CTX-M-15}}$, *bla*$_{\text{SHV-28}}$ (1) | *bla*$_{\text{CTX-M-15}}$, *bla*$_{\text{SHV-28}}$ (4)<br>*bla*$_{\text{SHV-28}}$ (1) | |
| | | KL116(1) | Col440I; IncA/C2; IncFIB(K); IncFII (1) | | *bla*$_{\text{NDM-1}}$, *bla*$_{\text{CTX-M-15}}$, *bla*$_{\text{SHV-28}}$ (1) | | | |

| | | | | | | | | |
|---|---|---|---|---|---|---|---|---|
| | | KL103(1) | Col440I; Col440II; IncFIB(K); IncFIB(pKPHS1); IncFIB(pQil); IncFII(K); IncFII(K); IncQ1; IncR (1) | $bla_{KPC-2}$ (1) | | | | |
| 219 (7) | O1v1(7) | KL114(7) | IncFIB(K) (5) | | | | $bla_{CTX-M-15}$ (1) | $bla_{CTX-M-15}$ (6) |
| 397 (7) | O1v1 (7) | KL158 (7) | FIA(pBK30683); IncFIB(K); IncFIB(Mar); IncFII(K); IncHI1B (4)<br>IncFIB(K); IncFIB(Mar); IncFII(K); IncHI1B (3) | | | | $bla_{CTX-M-15}$(7) | |
| 111 (6) | O1v2 (6) | KL63 (4) | FIA(pBK30683); IncFIB(K); IncFII(pKPX1) (2) | $bla_{NDM-1}$(3) | $bla_{NDM-1}$, $bla_{CTX-M-15}$(1) | | | |
| | | KL104 (2) | Col440I; IncA/C2; IncFIB(K); IncFII(K); IncFII(K) (2) | | | | $bla_{CTX-M-15}$(2) | |
| 307 (6) | O2v2 (6) | (KL102) | IncFIB(K)/IncFII(K); IncI1; IncQ1 (2) | | $bla_{NDM-1}$, $bla_{CTX-M-15}$, $bla_{SHV-28}$ (2) | | $bla_{CTX-M-15}$, $bla_{SHV-28}$ (1) | $bla_{SHV-28}$ (2), $bla_{CTX-M15}$, $bla_{SHV-28}$ (1) |
| 636 (6) | O2v1(4) | KL64(4) | IncA/C2; IncFIB(K); IncFII(K) (4) | $bla_{NDM-1}$(4) | | | | |
| | O5(2) | KL61(2) | IncA/C2; IncFIA(HI1); IncFIB(K); IncFII; IncFII(K) (2) | $bla_{NDM-1}$(1) | | $bla_{CTX-M-15}$ (1) | | |

**Supplementary Table 4.** Comparison of replicon-*bla*NDM gene combinations to reference plasmids and their distribution among STs and sentinel sites.

| Replicon (No. observed) | AMR Genes (No. observed) | No. of STs | No. of Sentinel Sites | No. of Isolates with >95% match | Reference Plasmid |
|---|---|---|---|---|---|
| IncFII(Yp) (48) | *bla*NDM-1, *rmtC*, *sul1* (46)<br>*bla*NDM-7, *rmtC*, *sul1* (1) | 14 | 6 | 47 | p13ARS_MMH0112-3 (Accession LR697124) |
| IncX3 (21) | *bla*NDM-7 (20) | 8 | 9 | 19 | p14ARS_MMH0055-5 (Accession LR697126) |

### SUPPLEMENTARY FIGURES

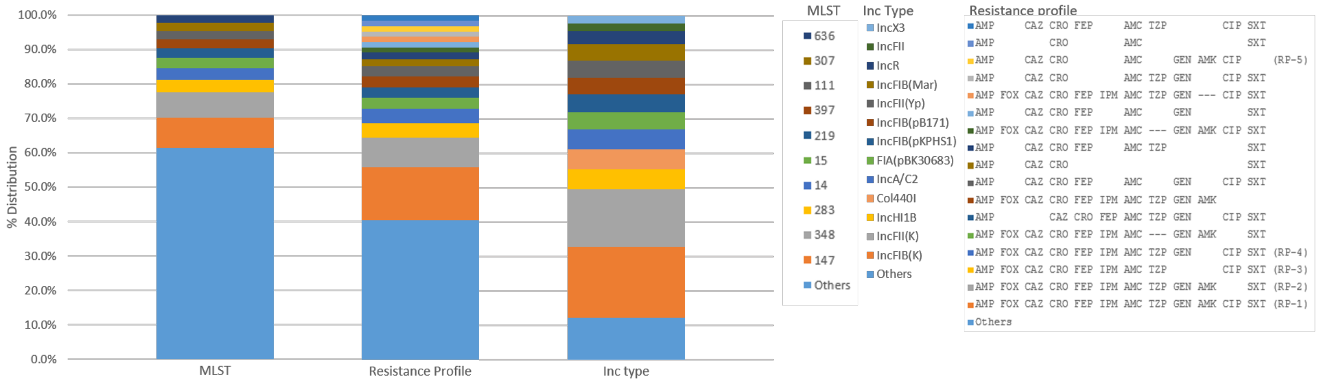


**Supplementary Figure 1.** Distribution of sequence types, resistance profiles (RP), and inc types among 259 *Klebsiella* isolates. The top 10 sequence types are shown while the remaining 94 are grouped as “Others.” The 10 most frequently observed inc type are shown while the remaining 22 are grouped as “Others.” The top 17 RPs are shown while the remaining 76 are grouped as “Others.”

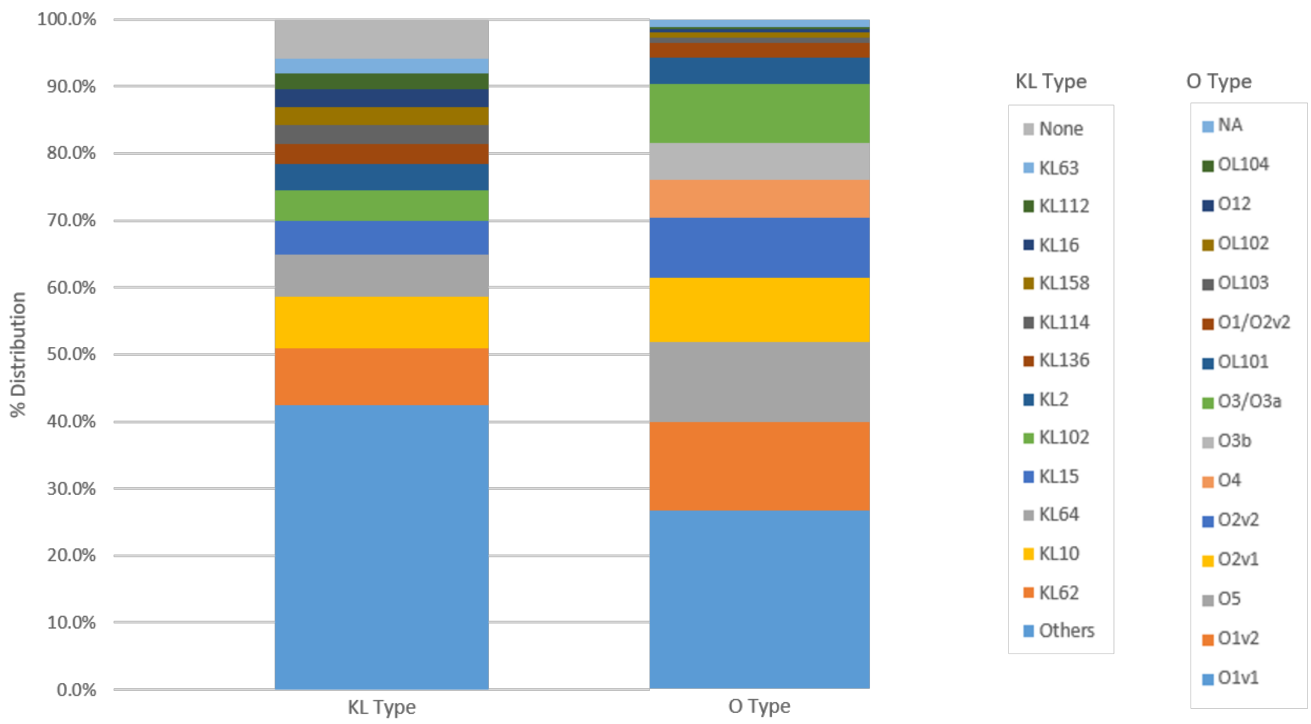


**Supplementary Figure 2.** Distribution of KL Types and O Locus Types among 259 *Klebsiella* isolates. The top 12 KL types are shown while the remaining 45 are grouped as "Others." All O locus types are shown with NA corresponding to O-types with low confidence scores.

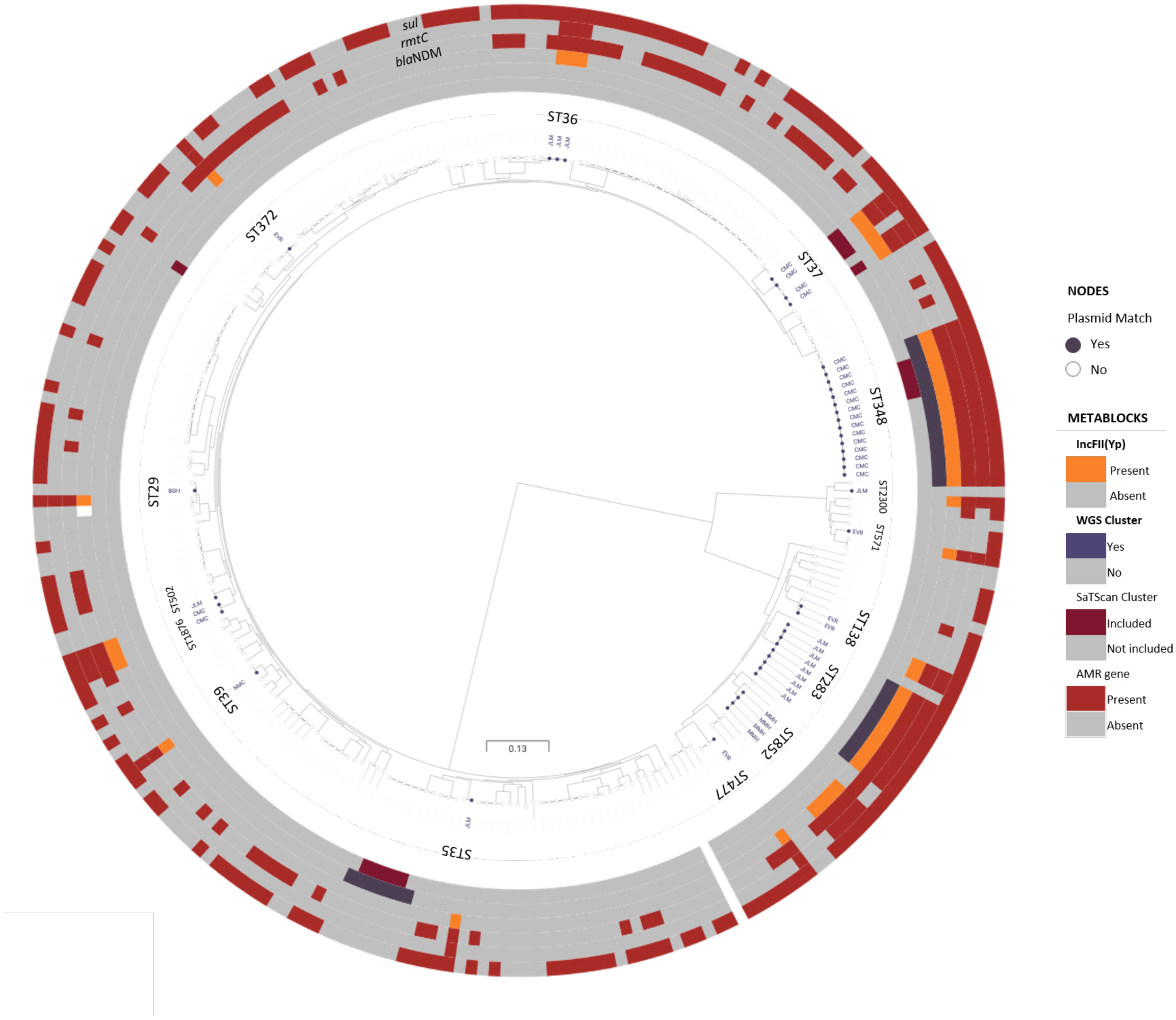


**Supplementary Figure 3.** Distribution of IncFII(Yp) replicon among the 259 *Klebsiella* isolates shows that CMC and JLM outbreaks isolates matched with >95% coverage to *bla*NDM-1-carrying plasmid p13ARS_MMH0112-3. The plasmid was also observed in various non-outbreak strains in CMC, JLM, and other sentinel sites.

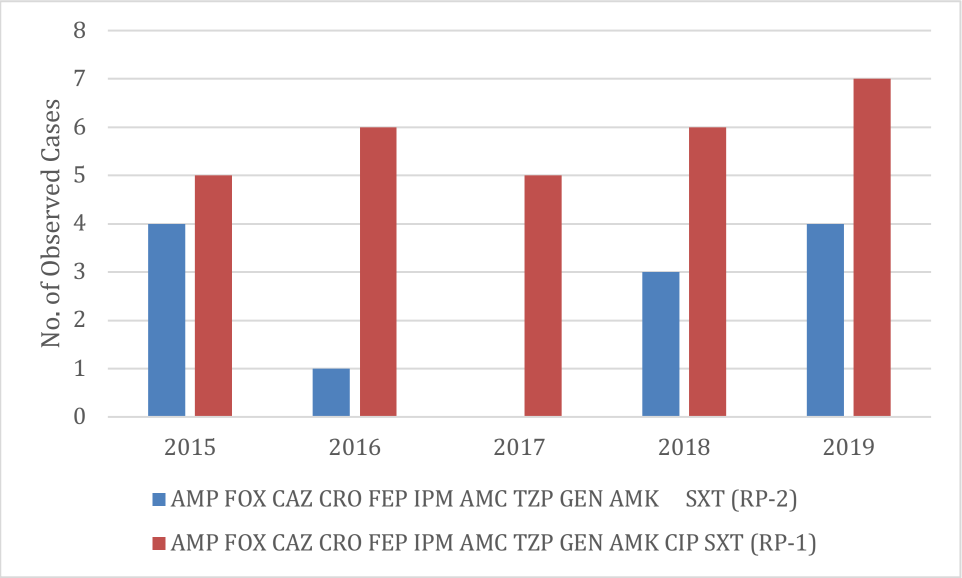


**Supplementary Figure 4.** Distribution of 41 *Klebsiella* isolates exhibiting RP-1 and RP-2 isolated in JLM from 2015-2019.

**SUPPLEMENTARY NOTE**

The members of the Department of Health Antimicrobial Resistance Surveillance Program are: Jerry Abrogueña, Christine Ivy Paula Agtuca, Maria Sarah Aguilar, Johari Ancheta, Evelyn M. Andamon, Myrna P. Angeles, Pauline Lois Bacungan, Ma. Teresa Barzaga, Maria Cecilia Belo, Marianette Noelle Bieren, Karen Bongas, Lucrecia Bongato, Julie Anne Cabantog, Donna Marie Calaoagan, Joebert Castillon, Frederick Castillon, Frederick Dalingding, Kristine dela Cruz, Joseph D. De Las Alas, Rowena Deloso, Floretess de Villa, Joselle Ealdama, Marie Karr Esguerra, Marilyn Espinosa, Arnold Joseph Fernandez, Hans Francis Ferraris, Racquel Florece, Joselyn Gacasan, Emil Bryan Garcia, Nelson Geraldino, Bernadette B. Hapitana, Louwela Jerusalem, Evelina Lagamayo, Modesty A. Leaño, Reynette Christine Ligaray, Ponciano Limcangco, Nena S. Lingayon, Sheryll Manzon, Ernesto Miralles, Ivan Ray Molina, Melissa Mondoy, Myra Olicia, Jane Pagaddu, Aireen Parayno, Grace Pong, Marniel Reyes, Chanda Romero, Annette Salillas, Jean Tan, Karlo Emir Tayzon, Elizabeth Freda Telan, Mary Ann Torregoza, Anacleta Valdez, Kristine Anne Vasquez, Ma. Merlina Vistal.

2. Clinical and Laboratory Standards Institute. Performance standards for antimicrobial susceptibility testing. 29th ed. Clinical and Laboratory Standards Institute Wayne, PA, 2019.
